# Multiple mechanisms regulate the nanoscale organization of PD-L1 at the cell surface

**DOI:** 10.64898/2026.08.31.748200

**Authors:** Guus A. Franken, Abbey B. Arp, Dora Čerina, Veerle van Esch, Blanca Scheijen, Annemiek B. van Spriel

## Abstract

The immune checkpoint protein PD-L1 plays a pivotal role in tumor immune evasion by binding to PD-1 on immune cells, including T lymphocytes. While the expression and function of PD-L1 have been well studied, the importance of its spatial organization on the cell surface of tumor cells remains poorly understood. In this study, we used super-resolution microscopy combined with biochemical perturbations to investigate the factors regulating PD-L1 clustering and its effects on PD-1 binding and T cell inhibition. We found that PD-L1 is organized into nanoscale clusters at the plasma membrane, with distinct regulatory roles for the actin cytoskeleton, galectin-3, and cholesterol. Disruption of cortical actin increased PD-L1 cluster size, while galectin-3 promoted smaller, denser clusters and increased PD-L1 lateral mobility. Cholesterol depletion reduced PD-L1 cluster size and number and impaired PD-1 binding. These findings indicate that PD-L1 surface organization is collectively regulated by the actin cytoskeleton, galectin-3, and membrane cholesterol within the plasma membrane of tumour cells. Our results provide new insights into the dynamic regulation of PD-L1 and its potential as a therapeutic target in cancer immunotherapy.

## Introduction

Programmed death-ligand 1 (PD-L1) is an immune checkpoint ligand that inhibits T cell responses by binding programmed cell death protein 1 (PD-1). Through upregulation of PD-L1 at the cell surface, tumor cells can evade immune surveillance. Antibody blockade therapy targeting PD-L1 or PD-1 can alleviate inhibition and can promote patient outcome in a range of different cancer types. However, therapy is not effective in all patients, which suggests additional levels of regulation ^1–3^.

Besides expression, the function of many cell surface proteins is regulated by their cell surface membrane organization which is often dynamic ^4,5^. Protein segregation into nano- or microscale membrane domains can promote functions such as signaling, antigen presentation, immune synapse formation, and cell migration. Major membrane organizers are galectins, the actin cytoskeleton, and lipid rafts. Galectins are soluble carbohydrate-binding proteins that recognize N-linked glycans with specific motifs and can crosslink glycoproteins to form clusters or affect their lateral mobility on the cell surface ^5–7^. The cortical actin cytoskeleton is located near the plasma membrane and is crucial in maintaining cell morphology and facilitating cell migration. Its mesh-like structure induces steric hindrance to some membrane proteins, which can lead to restricted membrane protein movement and protein clustering according to the ‘picket-fence’ model ^8^. Additionally, membrane proteins can be anchored to the cytoskeleton through linker proteins such as ezrin, radixin, and moesin (ERM) ^9^. Lipid rafts are proposed specialized nanosized membrane domains with a relatively high abundance of the sterol cholesterol and sphingolipids that consist of more saturated and longer fatty acids. These domains spontaneously form through liquid phase immiscibility, and preferentially contain membrane proteins with a longer and palmitoylated intracellular domain ^10,11^. Tetraspanins are a family of four-transmembrane proteins that can affect the surface expression, organization and function of interaction partners ^4,5^.

PD-L1 is a single-pass membrane protein predominantly located on the cell surface, which can traffick between the plasma membrane and intracellular compartments ^12–14^. The stability and turnover of PD-L1 is regulated through different post-translational modifications including N-linked glycosylation, palmitoylation, acetylation, and phosphorylation ^15–17^. In addition, glycosylation of PD-L1 is required for binding to PD-1 ^18^. Furthermore, PD-L1 surface expression is maintained by indirect interaction with the actin cytoskeleton through ezrin, radixin, and moesin (ERM) proteins ^19,20^. Additionally, PD-L1 contains cholesterol binding sites that stabilize surface expression ^21^. Moreover, galectin-3 binds to PD-L1, which both shields its antibody binding site and promotes PD-1 binding ^22,23^. In a giant plasma membrane vesicle (GPMV) model system, palmitoylation of PD-L1 enhances its localization to lipid rafts and this may promote a higher PD-1 affinity conformation of PD-L1 ^24^. Finally, we recently reported that the tetraspanin TSPAN4 interacts with PD-L1 and negatively affects PD-L1 cell surface expression and lateral mobility ^25^.

Although regulation of PD-L1 expression has been relatively well studied, how PD-L1 is spatially organized on the cell surface and how this affects PD-L1 functioning is not well understood. Here, we explored how PD-L1 is organized on the cell surface using super-resolution microscopy, and through multiple biochemical perturbations we define mechanisms of PD-L1 organization. We also studied how PD-L1 organization may influence PD-1 binding and T cell inhibition.

## Materials and methods

### Cell culture

BLM cells were maintained in Dulbecco’s Modified Eagle Medium (DMEM) (ThermoFisher Scientific, #31966-021) supplemented with 10% FBS (Cytiva, #SV30160.03) and 1% antibiotic-antimycotic (Gibco, #15240062) (DMEM CM). Jurkat T cells and SUP-HD1 cells were maintained in Roswell Park Memorial Institute (RPMI) 1640 Medium (ThermoFisher Scientific, #42401-018) supplemented with 1 % stable glutamine (Capricorn, #STA-B), 10% FBS and 1% antibiotic-antimycotic (Gibco, #15240062) (RPMI CM). Cells were cultured in a humidified incubator at 37 °C and 5% CO2, and were regularly checked for mycoplasma contamination.

### Flow cytometry

BLM cells were detached with 1.5 mM EDTA in PBS for 5 min at 37 °C, harvested and plated in v-bottom FACS plates, and stained with primary antibodies or recombinant human PD-1-Fc chimera protein (ThermoFisher Scientific, #A42530) and secondary antibodies for 30 min at 4 °C in PBA (PBS+ 1 % BSA + 0.05 % sodium azide). Staining with LIVE/DEAD™ Fixable Aqua Dead Cell Stain Kit (Thermofisher, #L34957) for 15 min at 4 °C was used for discerning live from dead cells. For intracellular staining and prior to filipin staining (Sigma-Aldrich, #SAE0088-1ML, 1:10), cells were fixed with 4 % paraformaldehyde (PFA) in PBS for 20 min at RT and quenched with 50 mM glycine (ThermoFisher Scientific, #ICN808831) in PBS for 10 min at RT. For permeabilization, cells were resuspended and further incubated with permeabilization buffer (Invitrogen, #00-5523-00).

Primary antibodies used: mouse anti PD-L1 for extracellular staining (ThermoFisher Scientific, #14-5983-82, 1:50), rat anti galectin-3 (Invitrogen, #14-5301-82, 1:25), rabbit anti PD-L1 for intracellular staining (Cell signaling technology, #13684S, 1:500), mouse IgG1 isotype (BioLegend, #400102, 1:50), rat IgG2a isotype (Invitrogen, #02-9688). Secondary antibodies used: goat anti mouse IgG1-Alexa-647 (ThermoFisher Scientific, #A21240, 1:400), goat anti rat IgG-Alexa-488 (ThermoFisher Scientific, # A11006, 1:400), goat anti human IgG-Alexa-488 (ThermoFisher Scientific, # A11013, 1:400), donkey anti-rabbit IgG F(ab’)₂ Fragment-PE (Jackson ImmunoResearch, #711-116-152, 1:200). Analysis was done with FACSVerse or FACSLyric flow cytometers (BD Biosciences).

### Quantification of total cell surface PD-L1 molecules

The total number of PD-L1 molecules at the cell surface of BLM WT cells was quantified using an indirect flow cytometry-based immunofluorescence assay (QIFIKIT®, BioCytex) following the manufacturer’s instructions. Primary mouse anti PD-L1 (ThermoFisher Scientific, #14-5983-82, 1:50) antibody was used.

### Plasmids

The plasmids PD-L1-FLAG and PD-1-FLAG were obtained from Genescript, and PX459 Cas9 vector was obtained from Addgene. Mutations in PD-L1 post-translational mutation sites and in cholesterol binding motifs were made using the Q5 Site-Directed Mutagenesis Kit (#E0554S, New England Biolabs) according to the manufacturer’s protocols and using custom-designed primers (Sigma-Aldrich). All constructs were verified by Sanger sequencing.

### BLM PD-L1 KO, PD-L2 KO, and PD-L1+PD-L2 double KO generation

BLM WT cells were transfected with PX459 Cas9 vector (Addgene) containing a guide RNA sequence targeting PD-L1 (oligo sense: CACCGCATAGTAGCTACAGACAGA, oligo anti-sense: AAACTCTGTCTGTAGCTACTATGC) which were obtained from Sigma-Aldrich and designed using CRISPOR.org. One day after transfection, untransfected cells were excluded by 1 µg/ml puromycin treatment for two days, after which cells were limited diluted and grown from monoclonal colonies. Knockout was validated by flow cytometry staining and genomic DNA isolation (Qiagen) followed by PCR of gene of interest using primers (Sigma-Aldrich) and Sanger sequencing. PD-L1+PD-L2 double KO and PD-L2 KO cells were obtained by transfecting monoclonal PD-L1 KO cells and WT cells respectively with PX459 Cas9 vector containing a guide RNA sequence targeting PD-L2 (oligo sense: CACCGCCAGGCTCAACATTAGCAGG, oligo anti-sense: AAACCCTGCTAATGTTGAGCCTGGC) which was obtained from Sigma-Aldrich and designed using CRISPOR.org. After 1 µg/ml puromycin treatment for two days, a polyclonal PD-L2 KO population was obtained by cell sorting with FACSMelody (BD Biosciences) after fluorescently staining cell surface PD-L2 and gating on PD-L2 negative cells. Knockout was validated by flow cytometry staining.

### Treatment with cytochalasin D

BLM WT cells were treated with 1.25 µg/ml cytochalasin D (cytD, Sigma-Aldrich, #C8273) or with dimethyl sulfoxide (DMSO) control in DMEM CM without FBS for 30 min at 37 °C. Cells were washed once with PBS and subsequently fixed (for IF, extracellular PD-L1 staining), or detached (for flow cytometry, extracellular PD-L1 staining).

### Treatment with recombinant galectin-3 and TD-139

BLM WT cells were treated with 1 µg/ml recombinant galectin-3 (Bio-Techne, #1154-GA), 50 µM galectin-3 inhibitor TD-139 (Olitigaltin, Selleck Europe, #S0471), or DMSO control in DMEM CM without FBS for 30 min at 37 °C. Cells were washed once with PBS and subsequently fixed (for IF, intracellular PD-L1 staining), or detached (for flow cytometry, extracellular PD-L1 staining).

### Treatment with methyl-β-cyclodextrin

BLM WT or PD-L2 KO cells were treated with several concentrations of methyl-β-cyclodextrin (MBCD) in DMEM CM without FBS for 30 min at 37 °C. Cells were washed once with PBS and subsequently fixed (for IF, intracellular PD-L1 staining), or detached (for flow cytometry, extracellular PD-L1 staining).

### Immunofluorescence microscopy

Cells were harvested with 1.5 mM EDTA in PBS for 5 min at 37 °C, counted, and 4x10^4^ cells were seeded on sterile 12 mm #1.5 thickness German glass cover slips (Electron Microscopy Sciences, # 72290-04) in a 24 well plate in DMEM CM. 24 hours after seeding, cells were treated with 50 ng/ml IFNγ. When appropriate, BLM PD-L1+PD-L2 KO cells were transfected by adding 50 µl Opti-mem containing 0.5 µg DNA and 2.4 µl PEI that was pre-incubated for 15 min at RT. After another 24 hours, following treatment or not, coverslips were washed 2x with PBS and fixed with 4 % PFA in PBS for 20 min at RT. Coverslips were blocked with blocking buffer (5 % BSA+0.3 M glycine+2 % human serum + 1 % goat serum in PBS) for 30 min at RT, and stained with primary and secondary antibodies in blocking buffer for 30 min at RT each, with 3x 5 min PBS washes in between. For intracellular staining, cells were permeabilized with 0.1% triton-X100 in PBS for 10 min at RT. Finally, cells were stained with 0.3 μg/ml 4’-6-diamidino-2-phenylindole (DAPI) for 2 min at RT, washed with PBS and milliQ, and embedded in Fluoromount G (ThermoFisher Scientific, # 00-4958-02). Imaging was performed using a Zeiss LSM900 confocal microscope equipped with an Airyscan Detector and a 63x Plan-Apochromat oil immersion 1.4 NA objective with Zeiss Zen software. Antibodies used: rabbit anti PD-L1 (intracellular domain binding, Cell signaling technology, #13684S, 1:500), mouse anti PD-L1 (extracellular domain binding, ThermoFisher Scientific, #14-5983-82, 1:50). For imaging F-actin, phalloidin-Alexa-488 (ThermoFisher Scientific, #A12379, 1:50) was used. Cholera toxin subunit B-Alexa-488 (Invitrogen, #C34775) was applied at 3 µg/ml in PBS for 30 min as a lipid raft probe. Secondary antibodies used: goat anti mouse IgG1-Alexa-647 (ThermoFisher Scientific, #A21240, 1:400), goat anti rabbit IgG-Alexa-647 (ThermoFisher Scientific, #A21245, 1:400), goat anti rabbit IgG-Alexa-568 (ThermoFisher Scientific, #A11036, 1:400).

### Direct stochastic optical reconstruction microscopy (dSTORM) microscopy

BLM WT cells were harvested with 1.5 mM EDTA in PBS for 5 min at 37 °C, counted, and 3x10^5^ cells were seeded on sterile 25 mm #1.5 thickness German glass cover slips (Electron Microscopy Sciences, #72290-12) in a 6 well plate in DMEM CM with 50 ng/ml IFNγ. After 24 hours, cells were fixed with 4% PFA +0.1 % glutaraldehyde in 0.2 M phosphate buffer pH 7.4 for 30 min at room temperature. For subsequent imaging, coverslips were washed with PBS and quenched with 100 mM glycine and 100 mM NH4Cl in PBS for 15 min at RT. Cells were then blocked with blocking buffer (BB, 50 mM glycine + 3 % BSA + 2 % human serum) for 1 hour and subsequently stained with mouse anti PD-L1 antibody (ThermoFisher Scientific, #14-5983-82, 1:50) in BB for 30 min at RT. After 2x 5 min washes in BB, cells were stained with an anti-mouse Alexa-647 conjugated nanobody (NanoTag Biotechnologies, #N2002, 1:100) in BB for 30 min at RT. After 2x 5 min washes in BB, cells were post-fixed with 4 % PFA in PBS for 15 min at RT. Coverslips were washed with PBS, mounted in OxEA buffer, and dSTORM acquisition was performed as described before ^26^.

### Co-immunoprecipitation

BLM WT cells were seeded at 2x10^6^ in T75 flasks 2-3 days before co-immunoprecipitation (co-IP). One day before co-IP, cells were treated with 50 ng/ml IFNγ. When appropriate, cells were additionally transfected with PD-L1-FLAG one day before co-IP experiments by adding 2 ml Opti-mem (ThermoFisher Scientific, #11058-021) to the cells containing 19.5 µg DNA and 94 µl PEI (Polysciences, #24765) that was pre-incubated for 15 min at RT.

Protein G sepharose beads (Cytiva, #17061801) were blocked with 3 % BSA or incubated with 10 µg/ml mouse IgG1 isotype antibody in PBS overnight. On the day of co-IP, subconfluent cells were detached with 1.5 mM EDTA in PBS for 5 min at 37 °C, harvested with PBS, and centrifuged at 413 g for 5 min before resuspension in DMEM CM containing 1 µg/ml r-gal-3 and incubation for 30 min at 37 °C. Then, cells were washed 3x with PBS, and lysed in ice-cold lysis buffer (1 % Brij97, 10 mM Tris-HCL (pH 7.5), 150 mM NaCl, 2 mM MgCl2, 2 mM CaCl2 in MQ, supplemented with protease inhibitor and phosphatase inhibitor) for 45 min with constant agitation at 4 °C. Insoluble material was removed by 6 min 3800 g centrifugation. Protein G sepharose beads were washed with lysis buffer, and pre-clear was performed with BSA-blocked and isotype-coated protein G sepharose beads for 1 hour at 4 °C with constant agitation. Thereafter, lysate was incubated with 3 µg antibody (rabbit anti FLAG (Merck, #F7425), mouse anti PD-L1 (ThermoFisher Scientific, #14-5983-82), or rat anti galectin-3 (Invitrogen, #14-5301-82)) for 1 hour at 4 °C with constant agitation, and subsequently with protein G sepharose beads for 2 hours at 4 °C with constant agitation. Afterwards, beads were washed 5x with ice-cold wash buffer (0.1 % Brij97, 10 mM Tris-HCl (pH 7.5), 150 mM NaCl, 2 mM MgCl2, 2 mM CaCl2 in MQ, supplemented with protease inhibitor and phosphatase inhibitor). Proteins were eluted from beads using 2x reducing laemmli sample buffer at 95 °C for 10 min and stored at -20 °C.

### Western blotting

For making whole cell lysates for western blotting, 7x10^5^ BLM PD-L1+PD-L2 KO cells were seeded in 6 well plate wells two days in advance. The next day, cells were transfected by adding 250 µl Opti-mem containing 2.5 µg DNA and 12 µl PEI that was pre-incubated for 15 min at RT. Total cell lysates were prepared by spinning down 1x10^6^ cells in PBS, resuspending in 1x laemmli buffer containing 2.5% β-mercaptoethanol (Sigma-Aldrich, #M3148) and leaving the lysate on ice for 10 min. Benzonase Nuclease (Sigma-Aldrich, #E1014) was added to reduce viscosity. Then, samples were boiled at 95 °C for 10 min and stored at -20 °C.

Protein samples were separated on a 12% SDS polyacrylamide gel and transferred to a PVDF membrane (GE Healthcare). The blot was blocked with Licor TBS or 5% BSA in PBS for 1 h at RT and incubated with primary antibodies (rabbit anti PD-L1 (Cell signaling technology, #13684S, 1:1000), rat anti galectin-3 (Invitrogen, #14-5301-82, 1:500), rabbit anti FLAG (Merck, #F7425, 1:1000), mouse anti vinculin (Sigma-Aldrich, #V9131-100UL, 1:1000)) overnight. After incubation with secondary antibody for 1 h at RT, the blot was scanned using Amersham Typhoon ™ (Cytiva). Washes were performed with TBS+0.05 % Tween-20. Blots were analyzed using Image Studio Light (LlCORbio). Secondary antibodies used: IRDye® 800CW Goat anti-Rabbit IgG (LICORbio, # 925-32211), IRDye® 680 Goat anti-Mouse IgG (LICORbio, # 926-32220), Alexa-680 Goat anti-Rat IgG (H+L) (ThermoFisher Scientific, #A-21096).

### Fluorescence recovery after photobleaching

BLM cells were seeded at 4x10^5^ cells in Willco dishes in DMEM CM one day before fluorescent recovery after photobleaching (FRAP). On the day of FRAP, cells were washed with Leibovitz medium (ThermoFisher Scientific, #21083027) and incubated with 10 µg/ml anti PD-L1 Alexa-488 antibody (ThermoFisher Scientific, #53-5983-42, 1:100) in Leibovitz medium for 15 min at 37 °C, after which cells were washed 3x and incubated with Leibovitz medium. FRAP experiments were performed with a Leica TCS SP8 SMD microscope equipped with a 60× water 1.2 NA objective (Leica) and an argon-ion laser set to bleach with 100% power at the 488 nm wavelength. The fluorescence intensity in the bleach zone as well as the whole cell and background was measured to correct for photobleaching and background signal. Immobile and mobile fractions were calculated manually, and the recovery curve and speed of recovery (T-half) was determined using the easyFRAP web tool ^27^.

### Stable PD-1-overexpressing Jurkat T cell line generation

For generating Jurkat T cells stably expressing PD-1, 2x10^6^ Jurkat T cells ^28^ were transfected with 2 µg PD-1-FLAG DNA using Neon transfection system according to manufacturer’s instructions, and starting two days after transfection for a period of three weeks, cells were sorted with FACSMelody after fluorescently labeling cell surface PD-1 (mouse anti PD-1-PE (BD Biosciences, #557946, 1:20)) and gating on PD-1 positive cells in order to obtain a Jurkat T cell population stably expressing PD-1.

### Jurkat co-culture assays

BLM PD-L2 KO cells were seeded two days before the Jurkat co-culture assay at 2x10^4^ cells per well in 96 well plates with 50 ng/ml IFNγ. At the day of the assay, medium was replaced with DMEM CM without FBS containing 1 µg/ml gp100 peptide (JPT Peptide Technologies, #SP-MHCI-0084-2), incubated for 1 hour at 37 °C, and washed 3x with DMEM CM. Then, cells were treated with MBCD or r-gal-3 for 30 min at 37 °C. 10 µg/ml durvalumab (Imfinzi, Medimmune/AstraZeneca, provided by RadboudUMC pharmacy) or human IgG1 isotype antibody (BioLegend, #403501) was added 15 min after the start of treatment. Afterwards, cells were washed 2x with PBS and fixed using 0.5 % PFA in PBS for 30 min at RT. Subsequently the fixative was removed, cells were quenched with 100 mM glycine in PBS for 30 min at RT, and then washed and replaced with RPMI CM. Jurkat T cells were collected, counted, and resuspended in fresh RPMI CM containing 4 µg/ml CD28 antibody (InVivoMAb, # BE0248), and 2x10^5^ cells were added on top of the BLM cells and incubated at 37 °C. After 24 hours, supernatant was collected and IL-2 ELISA (ThermoFisher Scientific, #88-7025-88) was performed.

### Data analysis

Flow cytometry analysis was performed using FlowJo X Software (FlowJo LLC).

Pearson and manders correlation analyses were performed using the BIOP JACoP plugin in Fiji image analysis software.

Cluster identification and quantification were performed using a custom macro script in Fiji image analysis software. For each image, the workflow proceeded as follows: first, a representative region of interest (ROI) of fixed size was determined for each cell and was duplicated to create a working copy. Local intensity maxima were detected using the Find Maxima function with a suitable prominence value, generating a segmented particle map. In parallel, a duplicate of the original image was thresholded using the Otsu dark method with suitable threshold, sharpened, and converted to a binary mask. The segmented maxima image and the thresholded binary mask were combined using the Image Calculator (AND) function to ensure that only maxima within thresholded regions were retained. The combined result was subjected to particle analysis (Analyze Particles, size: 0–∞ pixels), measuring morphological and intensity-based features including cluster area, cluster number, and median cluster intensity.

Scatter plots of cluster size or cluster number versus total intensity for cells overexpressing PD-L1 constructs were generated using R based on cluster quantification analysis using Fiji image analysis software as described above. Polynomial regression models (linear to cubic) were compared using the Akaike Information Criterion, and the best-fitting model was applied consistently across constructs. Regression lines were plotted with 95% confidence intervals.

For generating histograms showing PD-L1 cluster size distributions, cluster quantification analysis using Fiji image analysis software was performed as described above. Then, cluster area measurements from all identified clusters were combined for multiple cells using R and plotted with a defined cluster size range and bin width.

For dSTORM data analysis, localisation data was extracted using ThunderSTORM processing in Fiji, and DBSCAN analysis (ε = 80 nm, minpts = 5) was performed with the RSMLM package in R.

To estimate the mean number of PD-L1 molecules per cluster, cluster counts obtained by airyscan confocal microscopy, clustered fraction obtained by dSTORM microscopy, and molecule number obtained by QIFIKIT analysis were combined as follows,

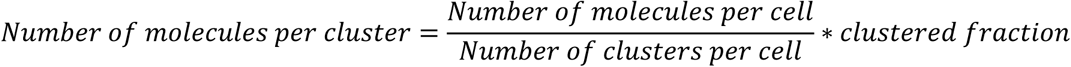

with number of clusters per cell defined as follows,

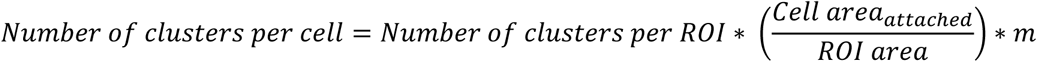

where *m* is defined as:

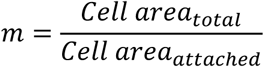

As *m*, a factor of 3 is taken, based on the assumption of adherent cells approximately resembling hemispheres.

### Statistical Analysis

Statistical analysis was performed using GraphPad Prism 9 (GraphPad). Normality was assessed using the Shapiro-Wilk test.

## Results

### PD-L1 is organized in nanoscale clusters on the plasma membrane of tumor cells

We examined PD-L1 endogenous expression on the cell surface of melanoma cell line BLM and Hodgkin lymphoma cell line SUP-HD1 by flow cytometry (Fig. 1A,B). We employed super-resolution airyscan confocal microscopy to investigate PD-L1 organization with ± 120 nm lateral resolution. Both melanoma and Hodgkin lymphoma cell lines showed PD-L1 organization in clusters that could be clearly defined by cluster analysis as ranging from 0.005 to 0.5 µm^2^ in size and with a median cluster size of ± 0.07 µm^2^ and 0.10 µm^2^ respectively, indicative of a nanoscale organization (Fig. 1C,D). Single molecule localization data obtained by super-resolution Direct Stochastic Optical Reconstruction Microscopy (dSTORM) confirmed strong clustering of PD-L1 with an average cluster size of ± 0.04 µm^2^ and that the majority of PD-L1 molecules resided in clusters with a clustered fraction of ± 0.90 (Fig. 1E-G). Interestingly, the PD-L1 clusters were similar in average size and distribution on these two different tumor cell lines. Quantitative analysis indicated that for BLM, an average cluster contained ± 8 PD-L1 molecules (Table 1). These results show PD-L1 is highly clustered in nanoscale domains on the surface of tumor cell lines as clearly defined structures.

**Figure 1:**
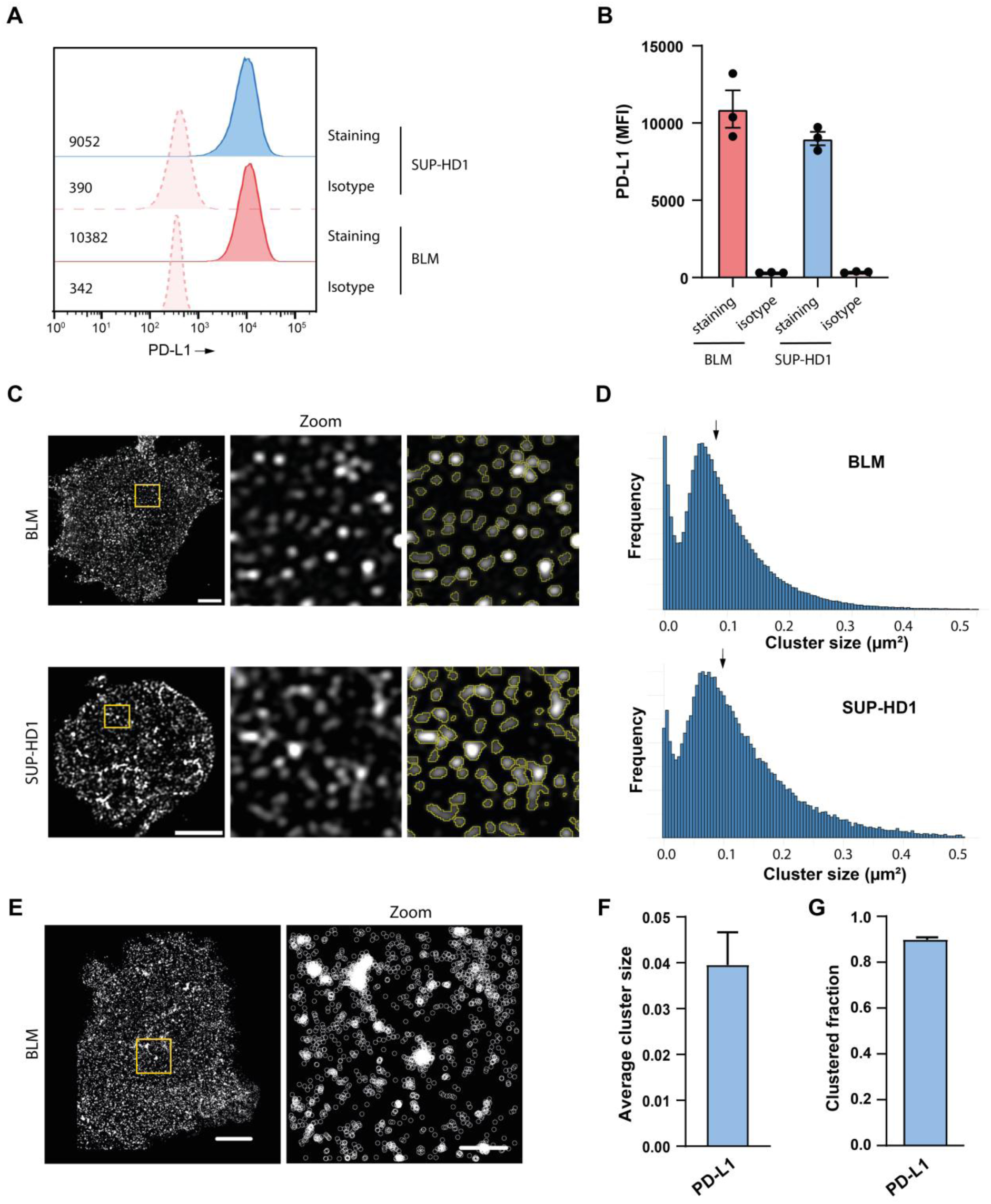
PD-L1 is organized in nanoscale clusters on the plasma membrane of tumor cells. (**A**) Representative flow cytometry histograms of PD-L1 surface expression on BLM (melanoma) and SUP-HD1 (Hodgkin lymphoma) cells from three independent experiments. (**B**) Flow cytometry surface PD-L1 median fluorescence intensity (MFI) values for BLM and SUP-HD1 cells for three independent experiments. Each dot represents an independent experiment. Data is shown as mean +/- SEM. (**C**) Representative immunofluorescence images of BLM and SUP-HD1 cells stained for PD-L1 obtained by airyscan confocal microscopy. Scale bar: 5 µm, for zoomed images 1 µm. (**D**) PD-L1 cluster size distribution with frequency from 438 pooled cells for BLM and 191 pooled cells for SUP-HD1 from one representative experiment. Arrows indicate median cluster size. (**E**) Representative dSTORM image of BLM cells stained for PD-L1. ROI in the left image is shown enlarged in the right image. Scale bar: 5 µm, for zoomed image 1 µm. (**F,G**) PD-L1 cluster size (F) and clustered fraction (G) on the cell surface of BLM cells as determined by dSTORM microscopy. Data is from 17 cells from two independent experiments. Data is shown as mean +/- SEM.

**Table 1:** PD-L1 cluster parameters in BLM cells.

| Number of clusters per cell | Number of molecules per cell | Clustered fraction | Number of molecules per cluster |
| --- | --- | --- | --- |
| 3645 ± 342 (n=47) | 33116 ± 3251 | 0.90 ± 0.043 (n=17) | 8.17 ± 1.11 |

Average number of PD-L1 molecules per cluster was estimated by dividing number of molecules per cell (obtained by QIFIKIT analysis) by number of clusters per cell (obtained by airyscan confocal microscopy), with a correction for the clustered molecular fraction (obtained by dSTORM microscopy). Data is shown as mean ± SEM from two independent experiments. When appropriate, number of measurements are indicated in brackets.

### The actin cytoskeleton restricts PD-L1 cluster size

PD-L1 expression is dependent on the actin cytoskeleton through interaction with ezrin, radixin, and moesin (ERM) proteins ^19,20^. To study the potential influence of cortical actin in PD-L1 organization, BLM cells were treated with cytochalasin D (cytD), which inhibits actin polymerization by binding to barbed ends of actin filaments ^29^. Treatment with cytD for 30 minutes did not change PD-L1 surface levels or viability of the tumor cells (Fig. 2A-C). CytD treatment altered a well-organized cortical actin network to fragmented and punctate actin structures, as indicated by phalloidin staining, confirming disruption of the actin cytoskeleton (Fig. 2D). Airyscan confocal microscopy and subsequent cluster analysis revealed that actin cytoskeleton disruption led to a small increase in cluster size (Fig. 2E,H), without affecting cluster number or intensity (Fig. 2F,G). These data indicate that the actin cytoskeleton limits PD-L1 cluster size.

**Figure 2:**
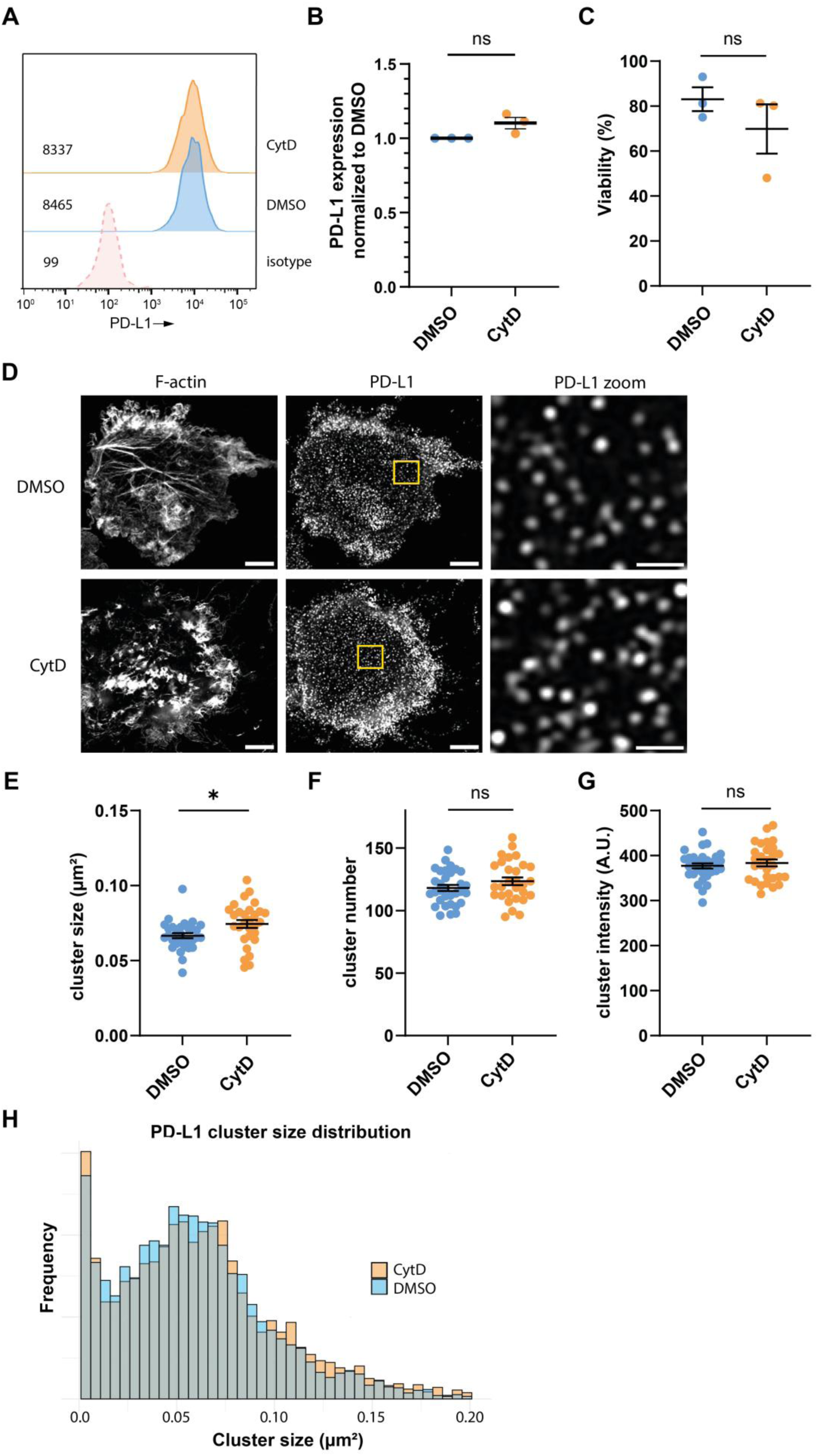
The actin cytoskeleton restricts PD-L1 cluster size. BLM cells were treated with cytochalasin D (cytD) at 1.25 µg/ml or DMSO for 30 min, before further processing. (**A**) Representative flow cytometry histograms of PD-L1 surface expression on BLM cells treated with cytD or DMSO out of three independent experiments. (**B,C**) Flow cytometry surface PD-L1 signal normalized to DMSO (B) and viability assessed by viability dye (C) for BLM cells treated with cytD or DMSO for three independent experiments. Each dot represents an independent experiment. Significance was determined by one sample Wilcoxon test (panel B, *P*=0.25) or Mann-Whitney test (panel C, *P*=0.70). (**D**) Representative immunofluorescence images of BLM cells stained for PD-L1 and F-actin after cytD or DMSO treatment obtained by airyscan confocal microscopy. Scale bar: 5 µm, for zoomed images 1 µm. (**E,F,G**) PD-L1 cluster size (E), cluster number (F), and cluster intensity (G) for BLM cells treated with cytD or DMSO for three independent experiments, obtained by airyscan confocal microscopy. All dots represent individual cells. Significance was determined by Mann-Whitney test (panel E, \**P*=0.0112) or unpaired t test (panel F, *P*=0.3076; panel G, *P*=0.7563). (**H**) PD-L1 cluster size distribution with frequency from 27 pooled cells for cytD-treated BLM cells and 32 pooled cells for DMSO treated BLM cells from one representative experiment. Data is shown as mean +/- SEM.

### Galectin-3 enhances PD-L1 cluster density and lateral mobility

Within the galectin family of carbohydrate-binding proteins, galectin-3 binds to PD-L1, shielding its antibody binding site, and promoting PD-1 binding ^22,23^. We investigated whether galectin-3 modulates PD-L1 organization. BLM cells that expressed moderate levels of galectin-3 were treated with recombinant galectin-3 (r-gal-3) or the galectin-3 inhibitor TD-139 ^30^, which did not affect PD-L1 surface expression or viability (Fig. 3A-C, Fig. S1A). Notably, r-Gal-3 treatment resulted in fewer and smaller-sized PD-L1 clusters, that were also more densely clustered as measured by cluster intensity (Fig. 3D-G). On the contrary, TD-139 treatment showed no impact on PD-L1 clustering, consistent with an opposing role of the inhibitor to galectin-3 (Fig. 3D-G, Fig. S1B).

**Figure 3:**
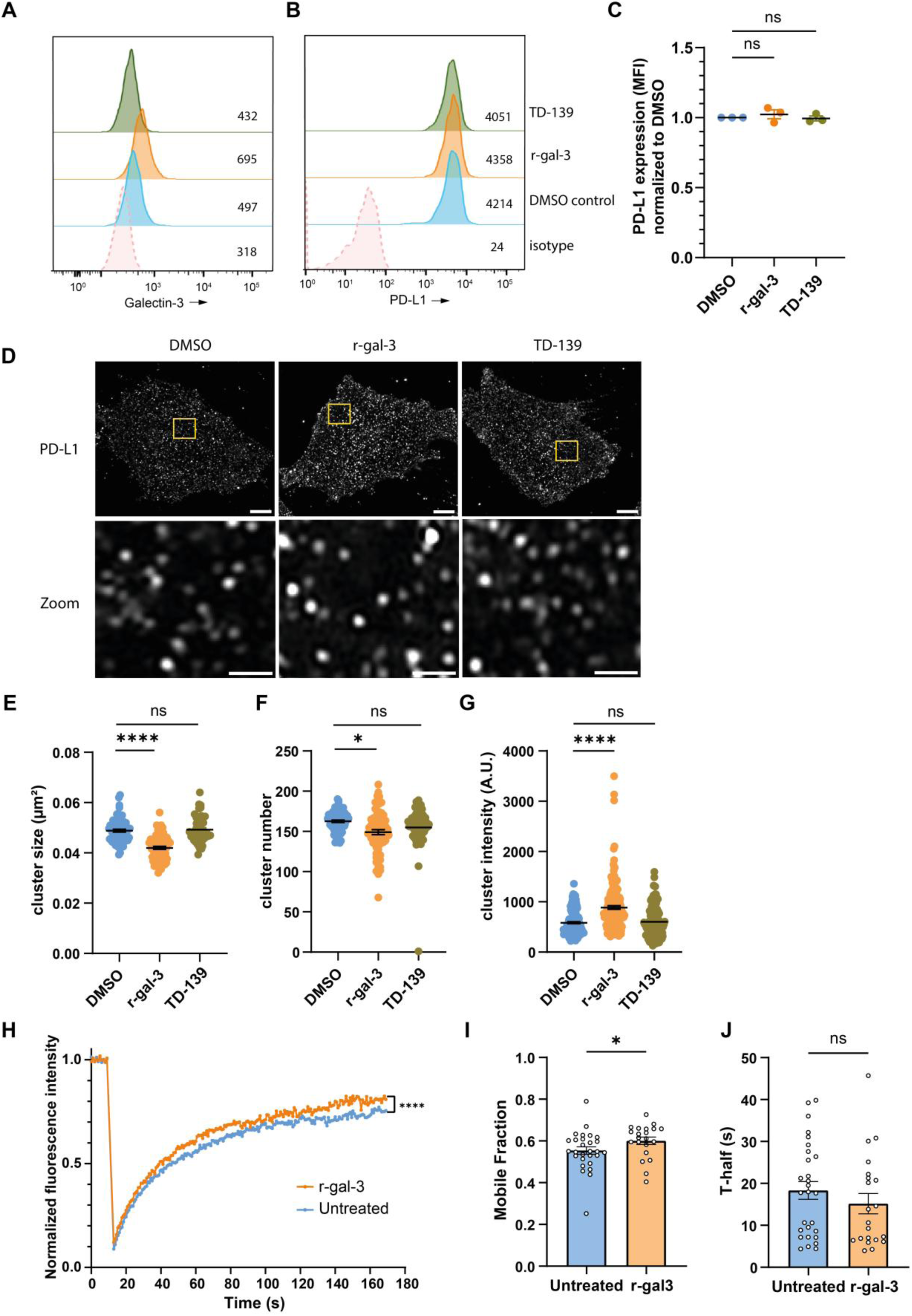
Galectin-3 enhances PD-L1 cluster density and lateral mobility. BLM cells were treated for 30 min with recombinant galectin-3 at 1 µg/ml, galectin-3 inhibitor TD-139 at 50 µM, or DMSO before further processing. (**A,B**) Representative flow cytometry histograms of galectin-3 (A) and PD-L1 (B) surface expression out of three independent experiments. (**C**) Flow cytometry surface PD-L1 signal of BLM cells for three independent experiments, normalized to DMSO. Each dot represents an independent experiment. Significance was determined by Wilcoxon signed rank test (ns, *P*>0.9999). (**D**) Representative immunofluorescence images of BLM cells stained for PD-L1 obtained by airyscan confocal microscopy. Scale bar: 5 µm, for zoomed images 1 µm. (**E,F,G**) PD-L1 cluster size (E), cluster number (F), and cluster intensity (G) for three independent experiments. All dots represent individual cells. Significance was determined by Kruskal-Wallis test with Dunn’s multiple correction (panel E: ns, *P*=0.6825,\*\*\*\**P*<0.0001; panel F: ns, *P*=0.3886,\**P*=0.0218; panel G: ns, *P*>0.9999,\*\*\*\**P*<0.0001). (**H**) Normalized fluorescence intensity recovery curves consisting of 21 recombinant galectin-3 treated and 28 untreated BLM cells from three independent experiments. Significance was determined by Wilcoxon matched-pairs signed rank test (\*\*\*\**P<*0.0001). (**I,J**) PD-L1 mobile fraction (I) and T-half (J) derived from three independent PD-L1 FRAP experiments. All dots represent individual cells. Significance was determined by Mann-Whitney test (panel I: \**P=*0.0274; panel J: ns, *P*=0.2736) Data is shown as mean +/- SEM.

Subsequently, we performed Fluorescence Recovery After Photobleaching (FRAP) on cell surface PD-L1 to investigate whether galectin-3 affected PD-L1 mobility. FRAP allows for the assessment of lateral dynamics of cell surface membrane proteins by tracking the rate and extent of fluorescence return following bleaching of a small region on the plasma membrane. Photobleaching of 5 µm spots on the plasma membrane revealed partial recovery of PD-L1 signal (Fig. 3H, Fig. S1C), with a mobile fraction of ± 55% (Fig. 3I) and a half-time of ± 18 seconds (Fig. 3J). These data indicate that a significant proportion of PD-L1 molecules at the cell surface exhibited restricted mobility. R-gal-3 increased the mobile fraction by ± 5–10% without substantially changing the half-time. Taken together, PD-L1 is laterally mobile on the surface of melanoma cells, and this mobility is increased by galectin-3.

To assess whether r-gal-3 might act directly on PD-L1 in modulating PD-L1 clustering and mobility, we performed co-immunoprecipitation experiments on r-gal-3 treated cells. However, no co-immunoprecipitation was observed between galectin-3 and PD-L1 (Fig. S1D), suggesting modulation of PD-L1 by galectin-3 to be indirect through interaction partners.

Since PD-L1 glycosylation is required for binding to PD-1 ^18^ and galectins typically bind specific glycosylated residues ^7^, we next investigated whether PD-L1 glycosylation is important in its plasma membrane organization. We generated PD-L1 KO BLM cells and transfected these cells with glycosylation site point mutated PD-L1 constructs showed that all four sites were indeed glycosylated, with N-linked glycosylation being the exclusive type of glycosylation as indicated by PNGase F treatment (Fig. S2A). Next, PD-L1 KO BLM cells were transfected with glycosylation site mutant constructs or a PD-L1 WT construct, stained for PD-L1 and imaged by airyscan confocal microscopy. Cluster size and number were plotted against total expression to account for expression differences between cells (Fig. S2B,C). We observed PD-L1 glycosylation site mutants did not show altered clustering as compared to wild-type PD-L1, both with respect to cluster size and cluster number. This indicates PD-L1 glycosylation as such does not impact PD-L1 plasma membrane organization. Therefore, these results indicate galectin-3 may promote PD-L1 cluster density and mobility, possibly through indirect mechanisms.

### Cholesterol is required for PD-L1 clustering and PD-1 binding

PD-L1 contains cholesterol binding sites that regulate the protein stability of PD-L1 ^21^. Therefore, we investigated the influence of plasma membrane cholesterol on PD-L1 organization in BLM cells. Methyl-β-cyclodextrin (MBCD) treatment was used to deplete plasma-membrane cholesterol ^11^, which did not affect PD-L1 expression levels or viability over a range of MBCD concentrations (Fig. 4A-C, Fig. S3A). Interestingly, PD-L1 cluster size and number, but not cluster intensity, were decreased by MBCD treatment (Fig. 4D-G, Fig. S3B).

**Figure 4:**
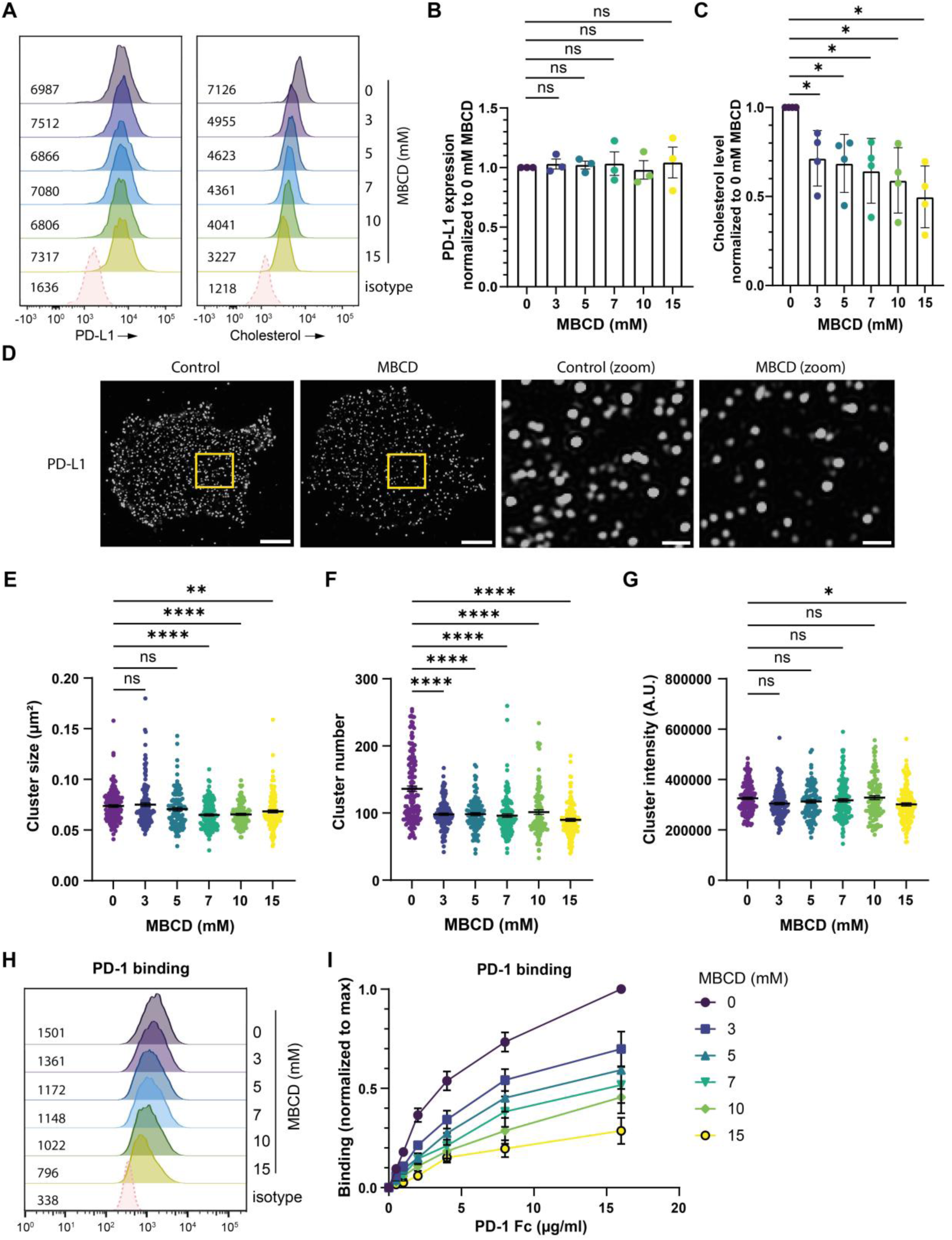
Cholesterol is required for PD-L1 clustering and PD-1 binding. BLM cells were treated for 30 min with several concentrations of MBCD before further processing. (**A**) Representative flow cytometry histograms of PD-L1 and cholesterol surface expression measured by filipin staining out of three independent experiments. (**B,C**) Flow cytometry surface PD-L1 (B) and cholesterol signal measured by filipin staining (C) for three or four independent experiments. Each dot represents an independent experiment. Significance was determined by one-sample Wilcoxon Signed Rank Test (panel B: ns, *P*=0.7500, *P*=0.7500, *P*>0.9999, *P*>0.9999, *P*=0.7500) or one-sample t test (panel C: * *P*=0.0351, * *P*=0.0308, * *P*=0.0298, * *P*=0.0209, * *P*=0.0103). (**D**) Representative immunofluorescence images of BLM cells stained for PD-L1 with or without 7 mM MBCD treatment, obtained by airyscan confocal microscopy. Scale bar: 5 µm, for zoomed images 1 µm. (**E,F,G**) PD-L1 cluster size (E), cluster number (F), and cluster intensity (G) for BLM cells treated with several concentrations of MBCD for three independent experiments, obtained by airyscan confocal microscopy. All dots represent individual cells. Significance was determined by Kruskal-Wallis test with Dunn’s multiple comparisons test (panel E: ns, *P*>0.9999, *P*>0.1048, \*\*\*\**P*<0.0001, \*\**P*=0.0056; panel F: \*\*\*\**P*<0.0001) or Brown-Forsythe and Welch ANOVA tests with Dunnett’s T3 multiple comparisons test (panel G: ns, *P*>0.9999, *P*=0.4526, *P*=0.8732, *P*=0.9996, \**P*=0.0115). (**H**) Representative flow cytometry PD-1 binding (16 µg/ml) histograms of cells treated with different concentrations of MBCD out of three independent experiments. (**I**) PD-1 binding to BLM PD-L2 KO cells at several concentrations following MBCD treatment for three independent experiments, normalized to the maximum signal per experiment. Data is shown as mean +/- SEM.

Lipid rafts are proposed nanoscale domains on the plasma membrane enriched in sphingolipids and cholesterol, and MBCD treatment impairs the stability of these domains ^11^. To gain mechanistic insight into the modulatory role of cholesterol, we investigated whether PD-L1 might be present in lipid rafts. However, PD-L1 did not colocalize to significant extents with the commonly used lipid raft probe cholera toxin subunit B ^11^ (Fig. S3C-E). This indicates that PD-L1 is most likely not present in lipid rafts, making PD-L1 organization by lipid rafts unlikely.

To determine whether cholesterol modulates PD-L1 organization through direct binding to PD-L1, we generated point mutations in PD-L1 that are reported to prevent cholesterol binding (F257A+R260A, F259A+R262A) ^21^. Furthermore, since PD-L1 palmitoylation has been proposed to alter its affinity to lipid rafts in a giant plasma membrane vesicle (GPMV) model system ^24^, we investigated the role of palmitoylation in PD-L1 organization by creating a point mutation in the PD-L1 palmitoylation site (C272A) ^15^. Whole cell lysates of transfected BLM PD-L1 KO cells showed comparable PD-L1 expression and indicated the mutants were unaltered in their glycosylation state compared to PD-L1 WT (Fig. S4A). Next, BLM PD-L1 KO cells were transfected with cholesterol binding site and palmitoylation site mutant expression plasmids or PD-L1 WT vector, stained for PD-L1 and imaged by airyscan confocal microscopy. Cluster size and number were plotted against total expression to account for expression differences between cells (Fig. S4B,C). All constructs showed similar PD-L1 cell surface clustering parameters compared to wild-type PD-L1. Together, these data indicate that direct binding of cholesterol to PD-L1 does not impact PD-L1 plasma membrane organization, and that cholesterol might indirectly modulate PD-L1 clustering.

To explore the functional consequence of PD-L1 organization in the context of PD-1 binding, we first generated programmed death-ligand 2 (PD-L2) KO BLM cells, since PD-L2 as a paralogous protein to PD-L1 also binds PD-1 ^31^. PD-L2 KO BLM cells were treated with MBCD and subsequently tested for recombinant PD-1 binding capacity. We observed that MBCD treatment impaired PD-1 binding in a concentration dependent manner (Fig. 4H,I). This indicates that PD-1 binding is affected by PD-L1 organization mediated by cholesterol.

To investigate this hypothesis, we generated PD-L1+PD-L2 KO BLM cells and transfected them with either the 4x PD-L1 cholesterol binding site mutant or with PD-L1 WT vectors and compared recombinant PD-1 binding capacity. For proper comparison, cells were gated on total PD-L1 to normalize for PD-L1 surface levels between constructs (Fig. S5A-D). Cells transfected with the 4x PD-L1 cholesterol binding site mutant did not differ in PD-1 binding at various concentrations as compared to cells transfected with PD-L1 WT (Fig. S5E,F). As a positive control, cells transfected with a PD-L1 construct with all four glycosylation sites mutated was impaired in PD-1 binding, consistent with earlier reports ^18,32^. These data suggest that the reported PD-L1 cholesterol binding sites do not impact PD-1 binding affinity. Together, these results suggest that cholesterol modulates both PD-L1 clustering and its affinity for PD-1 in an indirect manner.

### Galectin-3 and cholesterol do not impact PD-L1 mediated T cell inhibition

To determine whether galectin-3 or cholesterol-induced changes in PD-L1 organization affected PD-L1-mediated suppression of T cell activation, we used a gp100 peptide-dependent Jurkat T cell activation assay (Fig. 5A). BLM PD-L2 KO cells were treated with IFNγ, pulsed with gp100 peptide, treated with MBCD or r-gal-3, and fixed to preserve the organization state of PD-L1. Subsequently, Jurkat T cells expressing a gp100-specific T cell receptor, CD8 ^28^, and PD-1 were co-cultured overnight with the fixed cells, after which supernatants were analyzed for IL-2 to determine the Jurkat T cell activation state. Jurkat T cell activation was not induced without gp100 peptide with or without r-gal-3, excluding non-specific activation (Fig. 5B). Upon PD-L1 blockade using the therapeutic antibody durvalumab, IL-2 production was markedly increased compared to an isotype control antibody (Fig. 5B,D), confirming PD-L1-mediated inhibition. However, neither r-gal-3 nor MBCD treatment altered the degree to which PD-L1 inhibited Jurkat T cells (Fig. 5C,E). These data indicate that PD-L1 on BLM cells does not depend on galectin-3 or cholesterol to inhibit T cells through PD-1 binding.

**Figure 5:**
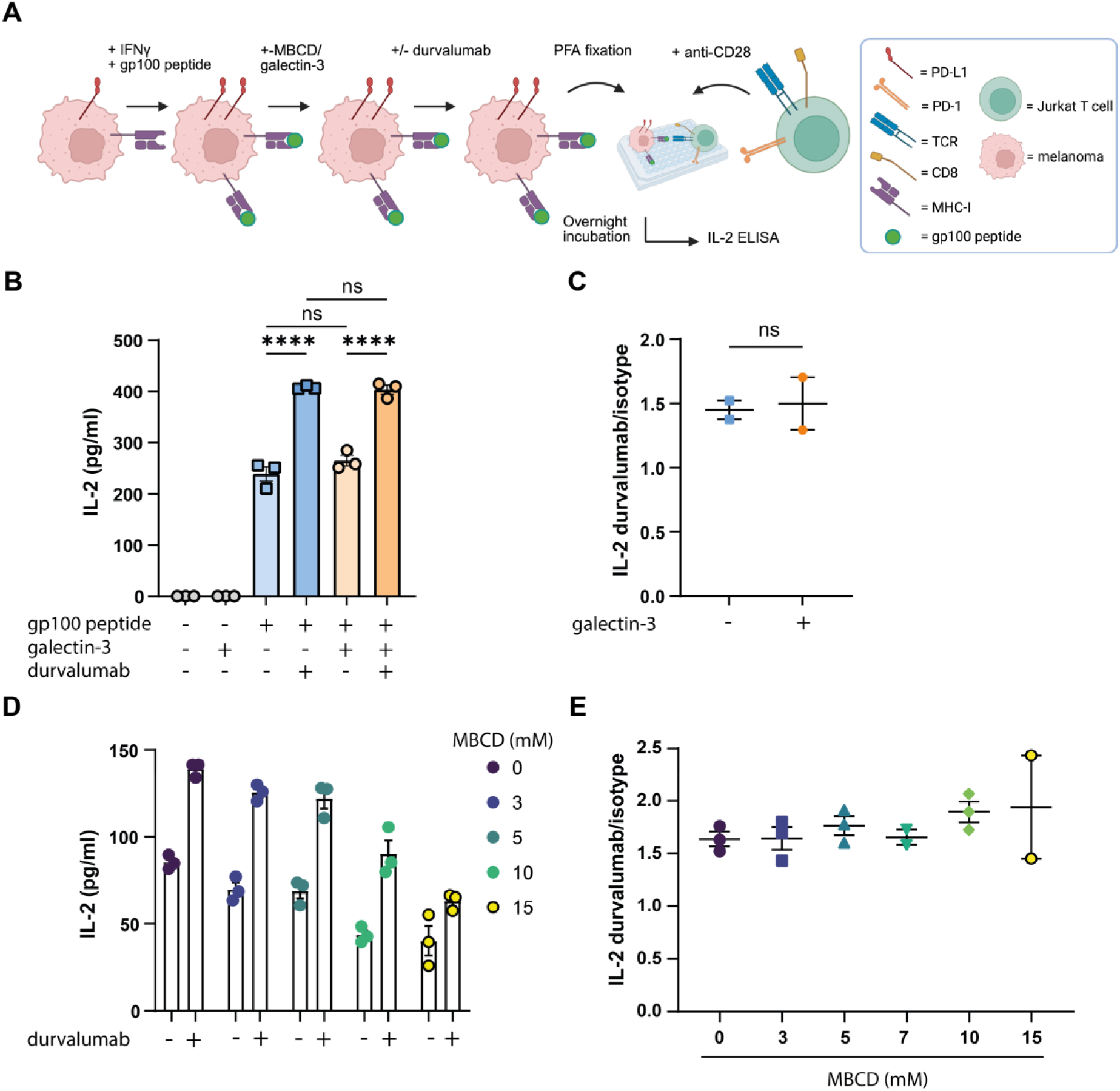
Galectin-3 and cholesterol do not impact PD-L1 mediated T cell inhibition. (**A**) Schematic of Jurkat-melanoma co-culture assay. BLM PD-L2 KO cells were treated with 50 ng/ml IFNγ for two days, peptide-pulsed with 1 µg/ml gp100 peptide for 1 hour, treated with MBCD at several concentrations or with recombinant galectin-3 at 1 µg/ml for 30 min, incubated with or without anti-PD-L1 (10 µg/ml durvalumab) for 20 min, and fixed and co-cultured with Jurkat T cells at a 1:4 BLM to Jurkat T cell ratio. After overnight incubation, supernatants were analyzed for IL-2 using ELISA. Schematic created with BioRender. (**B,D**) IL-2 production by Jurkat cells upon co-culture with BLM cells treated with recombinant galectin-3 (B) or MBCD (D) measured by ELISA. Dots are from n=3 technical replicates and are shown as mean +/- SEM. Significance was determined by one-way ANOVA with Tukey’s multiple comparisons test (panel B: ns, *P*=0.2804, *P*=0.9990, \*\*\*\**P*<0.0001). (**C,E**) IL-2 fold change of durvalumab-treated over isotype-treated samples for BLM cells treated with recombinant galectin-3 (C) or MBCD (E) upon co-culture with Jurkat T cells. Each dot represents an independent experiment. Significance was determined by paired t test (panel C: ns, *P*=0.7703).

## Discussion

PD-L1 antibody blockade therapy has proven to be effective in a wide range of cancer cell types, including metastatic melanoma and Hodgkin lymphoma ^1,2,33^. Many studies focus on regulation of PD-L1 expression, but not on PD-L1 surface organization. To better understand and predict differential outcome and potentially improve therapy, we examined whether PD-L1 function is influenced by its spatiotemporal membrane organization. Using super-resolution microscopy, we demonstrated that PD-L1 is highly organized in nanoscale clusters on the tumor cell surface, and that different organizational modulators have distinct effects. Since PD-L1 surface expression is maintained by interaction with the actin cytoskeleton through ERM proteins ^19,20^, we investigated its involvement in PD-L1 clustering. We showed that disturbing the actin cytoskeleton enlarges PD-L1 clusters. It might be that actin cytoskeleton perturbation releases ERM-restricted PD-L1 which is able to form larger clusters through other mechanisms by being allowed to freely diffuse. Galectin-3 regulates both PD-L1 affinity for PD-1 and blocking antibody efficacy ^22,23^. In contrast to the actin cytoskeleton, we showed galectin-3 has the ability to induce smaller, but more densely packed PD-L1 clusters.

In our system, PD-L1 glycosylation did not affect the degree of PD-L1 clustering on the cell surface. Since the most common method of galectin-3 binding to interaction partners is through binding glycosylation, these data suggest galectin-3 might influence PD-L1 clustering through modulating an interaction partner. Alternatively, galectin-3 might bind PD-L1 in a glycan-independent manner. Since the degree and type of glycosylation of PD-L1 at each glycosylation site is cell type specific, it remains to be seen whether glycosylation might play a role in PD-L1 organization in different cell types ^16,32^. Besides organization, galectin-3 treatment led to an increase in the PD-L1 mobile fraction. These data are in line with studies showing higher mobility upon recombinant galectin-3 treatment for α5β1 integrin on HeLa cells ^34^ and for N-cadherin in murine epithelial mammary tumor cells ^35^. Bigger or smaller clusters may be expected to behave differently in lateral mobility. Upon T cell recognition of antigen and proper co-stimulatory signaling, an immune synapse is formed which is characterized by protein organization in concentric domains ^5,36^. It would be interesting to see if galectin-3, in addition to promoting basal PD-L1 mobility, could affect PD-L1 recruitment to the immune synapse. In addition, single molecule tracking experiments can further provide insight about PD-L1 mobility.

Since cholesterol regulates PD-L1 expression through cholesterol binding domains, we investigated whether there is a role for cholesterol in PD-L1 membrane organization ^21^. Depleting plasma membrane cholesterol reduces PD-L1 cluster size and number, but does not affect cluster density. This suggests cholesterol promotes PD-L1 cluster formation by potentially recruiting unclustered or weakly clustered PD-L1 into larger assemblies. These findings suggest aspects of clustering such as cluster size and cluster density can be modulated independently of each other. It would be interesting to investigate the interplay between the factors that determine clustering. Employing single molecule localization microscopy techniques should be able to further elucidate how clustered and unclustered fractions change upon modulation of organization. Furthermore, high resolution live cell microscopy techniques could provide insight into the dynamics of PD-L1 clusters.

Cholesterol depletion affected PD-L1 organization, suggesting possible organization by lipid rafts. However, almost complete lack of PD-L1 colocalization with a lipid raft marker makes organization by lipid rafts unlikely. Furthermore, palmitoylation site or cholesterol binding site PD-L1 mutant constructs were not affected in organization. Although we confirmed the presence of glycosylation, we have not investigated to what extent palmitoylation and cholesterol binding actually take place for PD-L1 in our system, making it difficult to exclude these factors in PD-L1 organization. Our data suggest differential organization by cholesterol is mediated indirectly through an interaction partner. Since some tetraspanins are able to bind cholesterol and interference can lead to an altered organizing ability ^37^, it remains a possibility that cholesterol acts through tetraspanins in organizing PD-L1. We recently reported TSPAN4 as an interaction partner of PD-L1, which might have a possible role in organizing PD-L1.

PD-L1 does not dimerize or multimerize with itself ^38^, but it can interact *in cis* with a number of interaction partners, like CMTM and TRAPPC family members, integrin beta 4, and CD80 ^39–41^. Thus, in principle, our data does not exclude PD-L1 organization is mediated through the organization of interaction partners. In fact, our data regarding galectin-3 and cholesterol suggest these factors affect PD-L1 organization and mobility indirectly. Follow up experiments with for example *in vitro* membrane systems could allow for cluster observation in a more controlled environment.

We showed that upon cholesterol depletion the capacity for binding PD-1 was decreased, suggesting that a less organized PD-L1 state affects PD-1 affinity. This is in line with enhanced PD-1 binding observed upon artificially inducing PD-L1 tetramer clustering ^42^. Furthermore, a link between PD-L1 spatial organization and T cell inhibition was previously made based on DNA origami sheet assays ^43^. However, in our Jurkat T cell assay, the ability of PD-L1 in inhibiting T cells was unchanged upon cholesterol depletion of target cells. In these experiments, melanoma cells were fixed to preserve the PD-L1 organization and prevent endocytosis, in contrast to the PD-1 protein binding assay which was performed with live cells. Fixation may inadvertently impact membrane protein recognition, interfere with certain dynamic conformational changes between PD-L1 and PD-1 that is necessary for signal transduction ^44^, or interfere with proper immune synapse formation. Additionally, gp100 peptide concentration or PD-1 saturation might lead to suboptimal conditions. We also can not exclude that durvalumab binding affects PD-L1 organization, perhaps undoing any treatment effect of MBCD or r-gal-3. Future studies may benefit from blocking PD-L1 after fixation, ensuring no adverse effects on PD-L1 organization. Alternatively, *in vitro* membrane assays with artificially modulated PD-L1 organization might be able to shed more light on the importance of PD-L1 clustering on its function in inhibiting T cells, although this does not take the full complexity of living cells into account.

PD-L1/PD-1 antibody blockade therapy shows different degrees of efficacy between cancer types, with Hodgkin lymphoma having a much higher objective response rate than for example bladder cancer patients (87 compared to 24 % respectively) ^33^. It would therefore be interesting to investigate whether PD-L1 is differently organized between cancer cell types, whether shared organizational modulators exist, and whether this is related to clinical outcome.

## Conclusion

Our findings demonstrate that PD-L1 is predominantly organized in nanoscale clusters on tumor cell surfaces, with actin constraining cluster size, galectin-3 promoting smaller, denser clusters and lateral mobility, and membrane cholesterol supporting clustering and PD-1 binding. These multilayered organizational mechanisms modulate checkpoint engagement and may provide new avenues to predict and enhance immunotherapy efficacy.

## Abbreviations

BB: Blocking buffer
BSA: Bovine Serum Albumin
CM: Culture medium
co-IP: Co-immunoprecipitation
cytD: Cytochalasin D
DAPI: 4’-6-diamidino-2-phenylindole
dSTORM: Direct stochastic optical reconstruction microscopy
DMEM: Dulbecco’s Modified Eagle Medium
DMSO: Dimethyl sulfoxide
ELISA: Enzyme-Linked Immunosorbent Assay
ERM: Ezrin, radixin, and moesin
FACS: Fluorescence-activated cell sorting
FBS: Fetal Bovine Serum
FRAP: Fluorescence recovery after photobleaching
GPMV: Giant plasma membrane vesicle
IF: Immunofluorescence
IFNγ: Interferon-gamma
IgG: Immunoglobulin G
IL-2: Interleukin-2
KO: Knockout
MBCD: Methyl-β-cyclodextrin
PBA: PBS, BSA, and Sodium Azide solution
PBS: Phosphate-buffered saline
PCR: Polymerase Chain Reaction
PD-1: Programmed cell death protein 1
PD-L1: Programmed death-ligand 1
PD-L2: Programmed death-ligand 2
PEI: Polyethylenimine
PFA: Paraformaldehyde
PVDF: Polyvinylidene fluoride
r-gal-3: Recombinant galectin-3
ROI: Region of interest
RPMI: Roswell Park Memorial Institute medium
RT: Room temperature
SDS: Sodium dodecyl sulfate
TBS: Tris-buffered saline
WT: Wild-type

## Acknowledgements

We acknowledge funding support from the KWF Dutch Cancer Society (project 12949), the Netherlands Organization for Scientific Research ZonMW (project 09120012010023), and the European Research Council: Proof-of-Concept Grant (project 101112687). We thank the Radboudumc Microscopy Imaging Center for use of their microscopy facilities, as well as for their support and assistance.

## Author contributions

G.F.: Conceptualization; Formal analysis; Investigation; Visualization; Methodology; Writing-original draft; Writing-review and editing. A.A.: Investigation, Methodology, Formal analysis, and Visualization. D.C.: Formal analysis; Investigation; Visualization. V.E.: Formal analysis; Investigation; Visualization. B.S.: Supervision; Writing-review and editing. A.S.: Conceptualization; Supervision; Methodology; Writing-review and editing; Funding acquisition; Project administration.

