## Supplementary material for "Multiple mechanisms regulate the nanoscale organization of PD-L1 at the cell surface": Fig S1

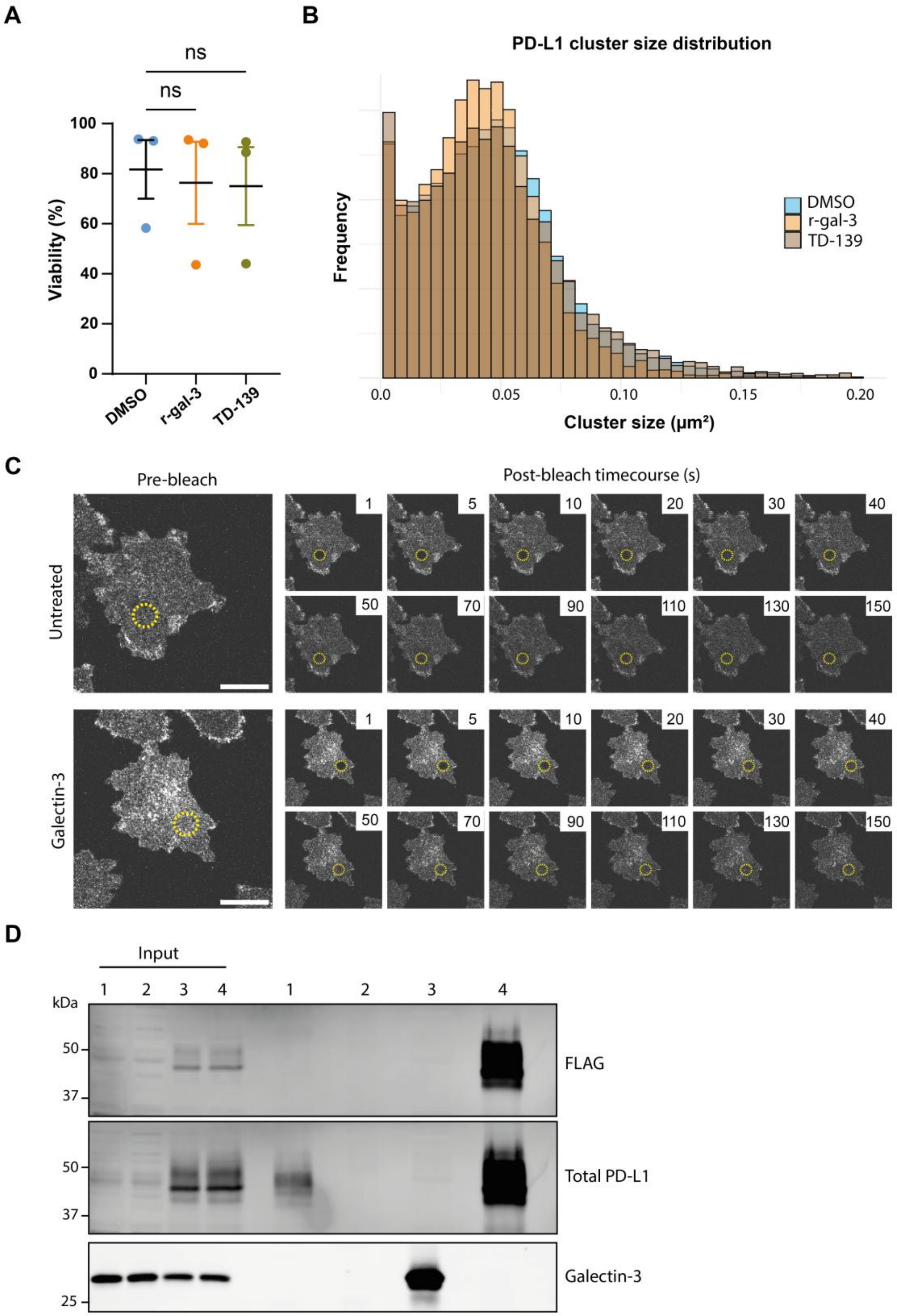


1. Untransfected --> PD-L1 IP
2. Untransfected --> isotype IP
3. PD-L1 FLAG transfected --> galectin-3 IP
4. PD-L1 FLAG transfected --> FLAG IP

**Figure S1:** Galectin-3 enhances PD-L1 cluster density and lateral mobility.

(**A**) Viability of BLM cells treated with recombinant galectin-3, TD-139, or DMSO for three independent experiments assessed by viability dye using flow cytometry. Each dot represents an independent experiment. Data is shown as mean +/- SEM. Significance is determined by Kruskal-Wallis test with Dunn's multiple comparisons test (ns, $P$=0.5934, $P$=0.9121). (**B**) PD-L1 cluster size distribution with frequency from 50 pooled recombinant galectin-3 treated BLM cells, 31 pooled TD-139 treated BLM cells, and 65 pooled DMSO treated BLM cells from one representative experiment. (**C**) Pre- and post-bleach images of a representative PD-L1 FRAP experiment with untreated or galectin-3 treated BLM cells. Circle indicates the area of bleaching and number indicates time after bleaching in seconds. Scale bar: 20 μm. (**D**) BLM cells were transfected with PD-L1 FLAG or not, and treated with r-gal-3 before being lysed. IP was performed on endogenous PD-L1, FLAG-tagged PD-L1, or galectin-3. Representative western blot showing FLAG signal, total PD-L1 signal, and galectin-3 signal.
