## Supplementary material for "Multiple mechanisms regulate the nanoscale organization of PD-L1 at the cell surface": Fig S2

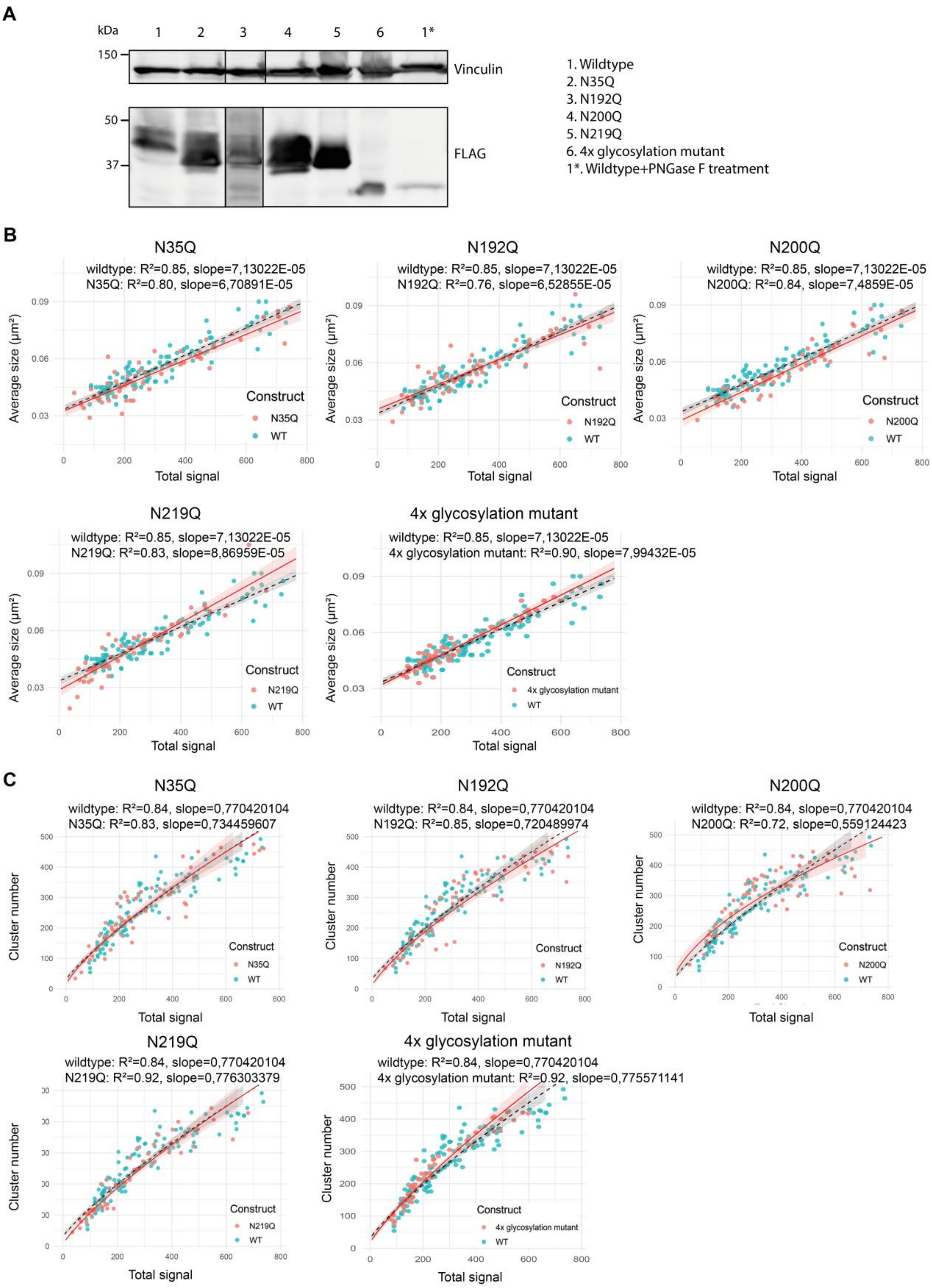
A
kDa
1 2 3 4 5 6 1*
150
50
37
Vinculin
FLAG
1. Wildtype
2. N35Q
3. N192Q
4. N200Q
5. N219Q
6. 4x glycosylation mutant
1*. Wildtype+PNGase F treatment
B
N35Q
wildtype: R²=0.85, slope=7,13022E-05
N35Q: R²=0.80, slope=6,70891E-05
N192Q
wildtype: R²=0.85, slope=7,13022E-05
N192Q: R²=0.76, slope=6,52855E-05
N200Q
wildtype: R²=0.85, slope=7,13022E-05
N200Q: R²=0.84, slope=7,4859E-05
N219Q
wildtype: R²=0.85, slope=7,13022E-05
N219Q: R²=0.83, slope=8,86959E-05
4x glycosylation mutant
wildtype: R²=0.85, slope=7,13022E-05
4x glycosylation mutant: R²=0.90, slope=7,99432E-05
Average size (µm²)
Total signal
Construct
WT
C
N35Q
wildtype: R²=0.84, slope=0,770420104
N35Q: R²=0.83, slope=0,734459607
N192Q
wildtype: R²=0.84, slope=0,770420104
N192Q: R²=0.85, slope=0,720489974
N200Q
wildtype: R²=0.84, slope=0,770420104
N200Q: R²=0.72, slope=0,559124423
N219Q
wildtype: R²=0.84, slope=0,770420104
N219Q: R²=0.92, slope=0,776303379
4x glycosylation mutant
wildtype: R²=0.84, slope=0,770420104
4x glycosylation mutant: R²=0.92, slope=0,775571141
Cluster number
Total signal
Construct
WT

**Figure S2:** The effect of glycosylation on PD-L1 clustering on the plasma membrane.

(**A**) Western blot showing molecular weight of PD-L1 glycosylation site mutants and wild-type PD-L1 after PNGase F treatment compared to wild-type PD-L1, obtained from transfected BLM PD-L1 KO cells and stained by FLAG tag. Vinculin is included as a loading control. Molecular weight marker is in kDa. (**B,C**) Average cluster size (B) and cluster number (C) plotted against total PD-L1 expression for PD-L1 glycosylation site mutants in transfected BLM PD-L1 KO cells, as obtained by airyscan confocal microscopy. All dots represent individual cells from two independent experiments. Curve parameters are indicated, together with a 95% shaded confidence interval.
