## Supplementary material for "Multiple mechanisms regulate the nanoscale organization of PD-L1 at the cell surface": Fig S3

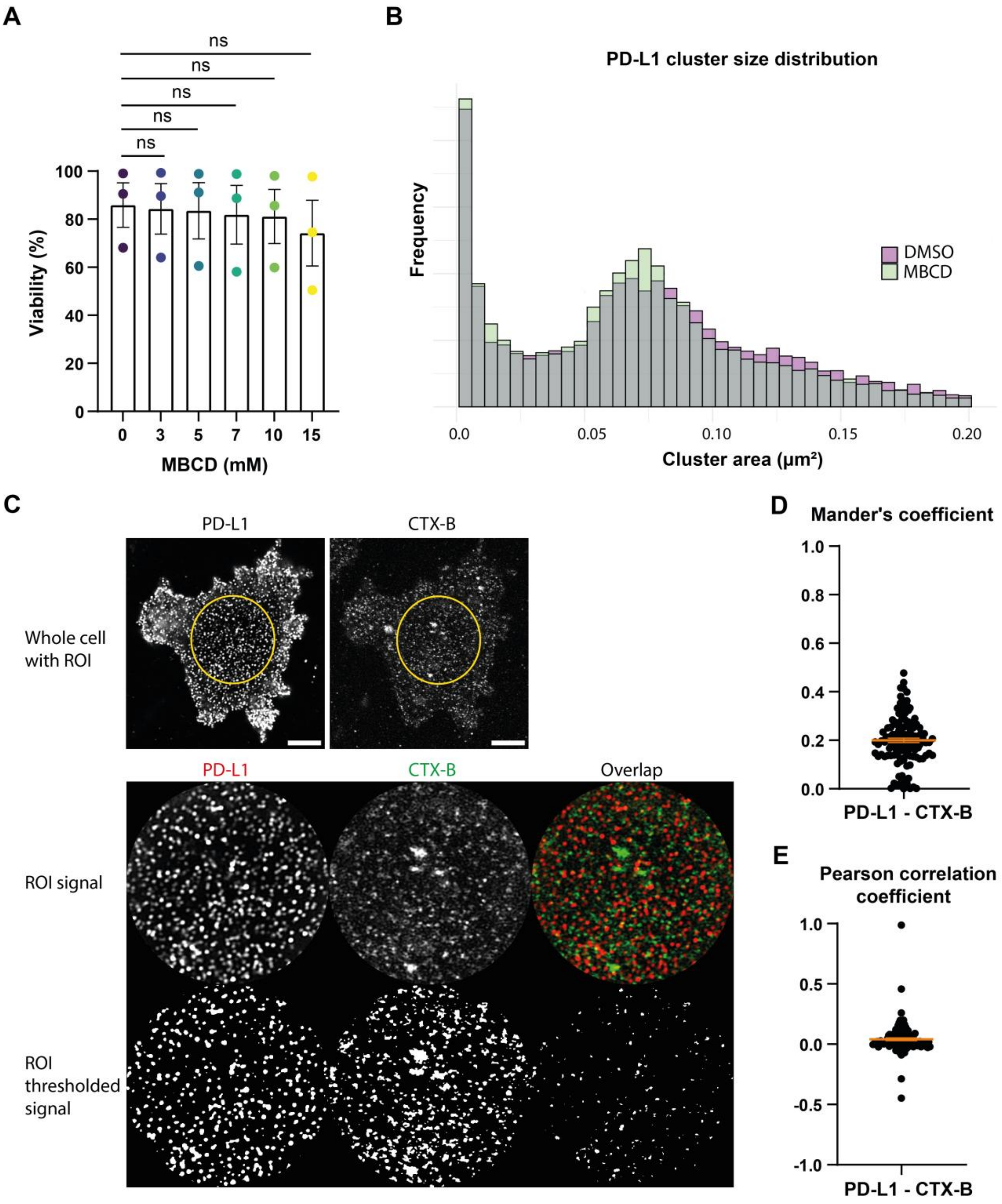
A
ns
ns
ns
ns
ns
100
80
60
40
20
0
Viability (%)
0
3
5
7
10
15
MBCD (mM)
B
PD-L1 cluster size distribution
Frequency
DMSO
MBCD
0.0
0.05
0.10
0.15
0.20
Cluster area (μm²)
C
PD-L1
CTX-B
Whole cell with ROI
PD-L1
CTX-B
Overlap
ROI signal
ROI thresholded signal
D
Mander's coefficient
1.0
0.8
0.6
0.4
0.2
0.0
PD-L1 - CTX-B
E
Pearson correlation coefficient
1.0
0.5
0.0
-0.5
-1.0
PD-L1 - CTX-B

**Figure S3:** Cholesterol is required for PD-L1 clustering and PD-L1 does not localize to lipid rafts.

(**A**) Viability of BLM cells treated with MBCD for three independent experiments assessed by viability dye using flow cytometry. Each dot represents an independent experiment. Data is shown as mean +/- SEM. Significance was determined by Kruskal-Wallis test with Dunn's multiple comparisons test (ns, $P$>0.9999). (**B**) PD-L1 cluster size distribution with frequency from 68 pooled cells for MBCD (7 mM) treated BLM cells and 61 pooled cells for DMSO treated BLM cells based on airyscan microscopy from one representative experiment. (**C**) Representative immunofluorescence images of BLM cells stained for PD-L1 and lipid raft marker cholera toxin subunit B (CTX-B) out of three independent experiments, obtained by airyscan confocal microscopy. Scale bar: 5 µm. Depicted is overlap between PD-L1 and CTX-B without and with a threshold applied to both PD-L1 and CTX-B signal. (**D,E**) Manders (PD-L1 in CTX-B) (D) and Pearson correlation coefficient (E) between PD-L1 and CTX-B, obtained by airyscan confocal
