## Supplementary material for "Multiple mechanisms regulate the nanoscale organization of PD-L1 at the cell surface": Fig S4

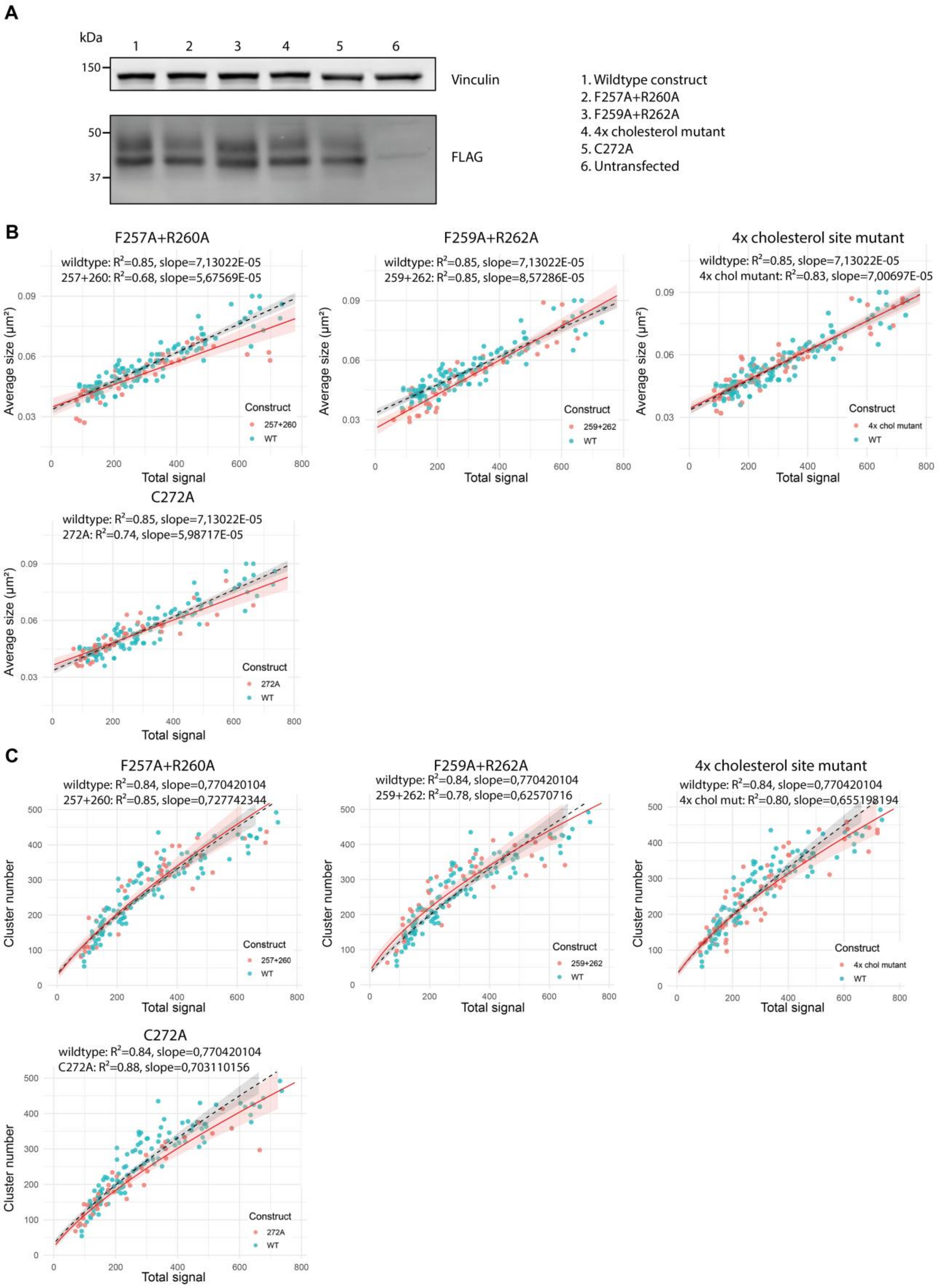
A
kDa
1
2
3
4
5
6
150
Vinculin
50
37
FLAG
1. Wildtype construct
2. F257A+R260A
3. F259A+R262A
4. 4x cholesterol mutant
5. C272A
6. Untransfected
B
F257A+R260A
wildtype: R²=0.85, slope=7,13022E-05
257+260: R²=0.68, slope=5,67569E-05
F259A+R262A
wildtype: R²=0.85, slope=7,13022E-05
259+262: R²=0.85, slope=8,57286E-05
4x cholesterol site mutant
wildtype: R²=0.85, slope=7,13022E-05
4x chol mutant: R²=0.83, slope=7,00697E-05
C272A
wildtype: R²=0.85, slope=7,13022E-05
272A: R²=0.74, slope=5,98717E-05
Average size (µm²)
Total signal
Construct
257+260
259+262
4x chol mutant
272A
WT
C
F257A+R260A
wildtype: R²=0.84, slope=0,770420104
257+260: R²=0.85, slope=0,727742344
F259A+R262A
wildtype: R²=0.84, slope=0,770420104
259+262: R²=0.78, slope=0,62570716
4x cholesterol site mutant
wildtype: R²=0.84, slope=0,770420104
4x chol mut: R²=0.80, slope=0,655198194
C272A
wildtype: R²=0.84, slope=0,770420104
C272A: R²=0.88, slope=0,703110156
Cluster number
Total signal
Construct
257+260
259+262
4x chol mutant
272A
WT

**Figure S4:** The effect of cholesterol binding sites and palmitoylation on PD-L1 clustering on the plasma membrane.

(**A**) Representative western blot out of three independent experiments of PD-L1 cholesterol binding site mutants and of PD-L1 palmitoylation (C272A) mutant compared to wild-type PD-L1, obtained from transfected BLM PD-L1 KO cells and stained by FLAG tag. Vinculin is included as a loading control. Molecular weight marker is in kDa. (**B,C**) Average cluster size (B) and cluster number (C) plotted against total PD-L1 expression for PD-L1 cholesterol binding site mutants and of the PD-L1 palmitoylation (C272A) mutant in transfected BLM PD-L1 KO cells, as obtained by airyscan microscopy. All dots represent individual cells from two independent experiments. Curve parameters are indicated, together with a 95% shaded confidence interval.
