## Supplementary material for "Multiple mechanisms regulate the nanoscale organization of PD-L1 at the cell surface": Fig S5

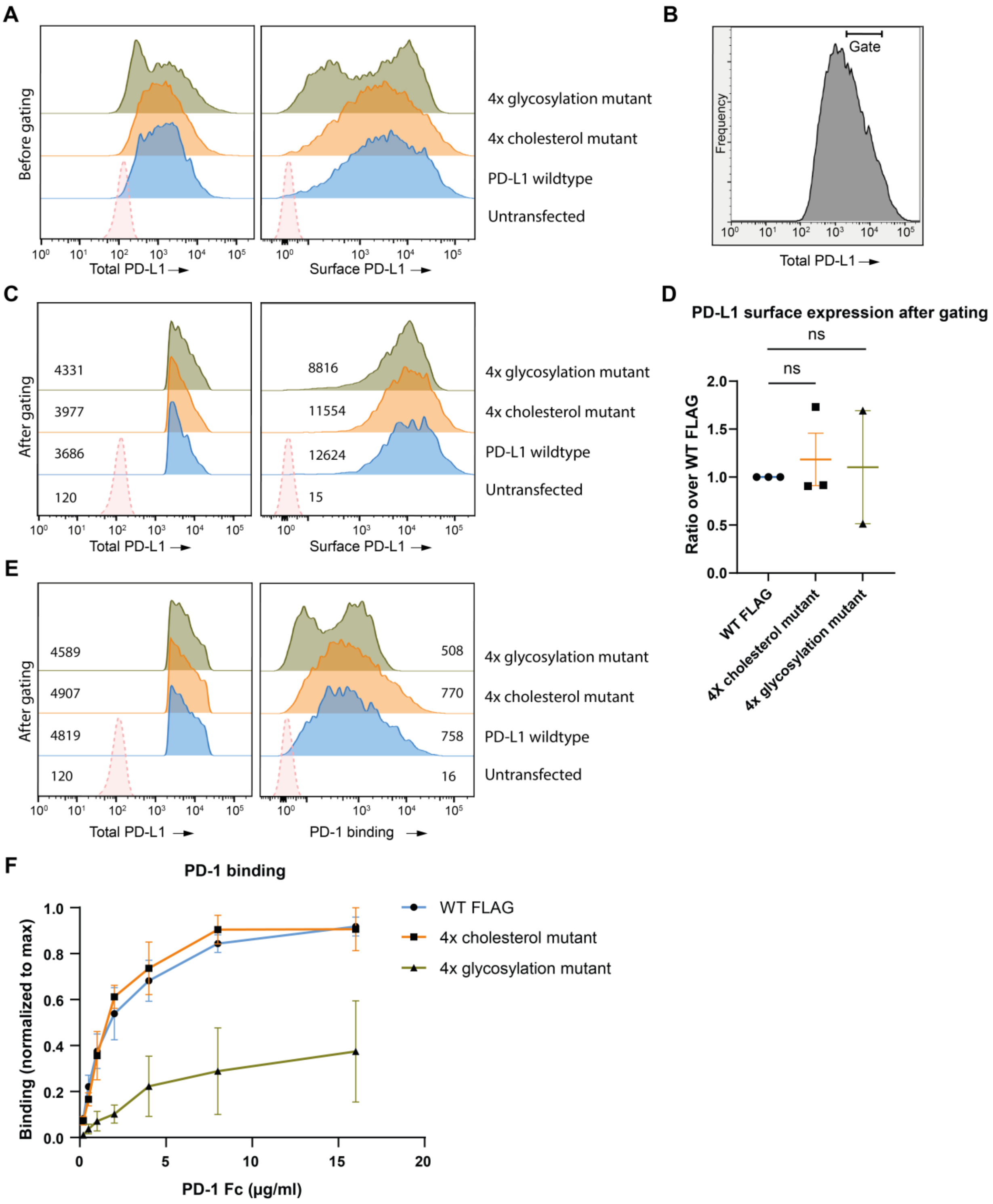
A
Before gating
Total PD-L1
Surface PD-L1
4x glycosylation mutant
4x cholesterol mutant
PD-L1 wildtype
Untransfected
B
Gate
Frequency
Total PD-L1
C
After gating
4331
3977
3686
120
8816
11554
12624
15
Total PD-L1
Surface PD-L1
4x glycosylation mutant
4x cholesterol mutant
PD-L1 wildtype
Untransfected
D
PD-L1 surface expression after gating
ns
ns
Ratio over WT FLAG
WT FLAG
4X cholesterol mutant
4x glycosylation mutant
E
After gating
4589
4907
4819
120
508
770
758
16
Total PD-L1
PD-1 binding
4x glycosylation mutant
4x cholesterol mutant
PD-L1 wildtype
Untransfected
F
PD-1 binding
Binding (normalized to max)
PD-1 Fc (μg/ml)
WT FLAG
4x cholesterol mutant
4x glycosylation mutant

**Figure S5:** Cholesterol binding sites do not affect PD-1 binding affinity of PD-L1.

BLM PD-L1+PD-L2 KO cells were transfected with PD-L1 wild-type, 4x glycosylation site mutant or 4x cholesterol binding site mutant constructs. (**A,B,C**) Representative flow cytometry total (using a PD-L1 C-terminal targeting antibody) and surface PD-L1 expression (using a PD-L1 N-terminal targeting antibody) histograms before (A) or after (C) gating applied to total PD-L1 (B). (**D**) PD-L1 surface expression after total PD-L1 gating as in (B) for BLM cells transfected with PD-L1 4x glycosylation site mutant and 4x cholesterol binding site mutant constructs relative to wild-type PD-L1 construct for three independent experiments. Each dot represents and independent experiment. Significance is determined by one sample t test (ns, $P$=0.5711, $P$=0.8900). (**E**) Representative flow cytometry total PD-L1 expression and PD-1 (2 μg/ml) binding histograms of PD-L1 wild-type, 4x glycosylation site mutant, and 4x cholesterol binding site mutant transfected cells after gating on total PD-L1 as in (B). (**F**) PD-1 binding to BLM cells transfected with PD-L1 wild-type, 4x glycosylation site mutant, and 4x cholesterol binding site mutant after gating on total PD-L1, normalized to highest signal per experiment for three independent experiments. Data is shown as mean +/- SEM.
